# Pushing for survival: Spatial intermixing and indirect resistance enable collective growth

**DOI:** 10.64898/2026.05.07.723406

**Authors:** Mireia Cordero, Brage Haldor Thomsen, Artemis Talliou, Anja K. Ehrmann, Sine Lo Svenningsen, Namiko Mitarai, Liselotte Jauffred

## Abstract

The survival of bacterial communities depends on complex dynamics at molecular, cellular, and ecosystem levels. Understanding antibiotic resistance requires a broader community context, as emergent dynamics can lead to unexpected outcomes, such as the persistence of susceptible populations or community collapse. We capture these behaviors by integration of microscopy and mathematical modeling to understand how bacterial interactions and spatial organization shape bacteriostatic antibiotic resistance in a two-strain community. We show that local chloramphenicol detoxification and mechanical pushing shape bacterial coexistence and spatial organization, promoting the survival and growth of otherwise susceptible bacteria. Additionally, the timing of antibiotic administration critically determines the growth dynamics, co-existence, local diversity of susceptible and resistant bacteria, and overall community resistance. Together, these insights highlight how community-level interactions fundamentally reshape antibiotic responses and open new avenues to understand and control bacterial resilience.

**SIGNIFICANCE STATEMENT:** Antibiotic resistance is usually treated as a property of individual bacterial strains, yet bacteria typically grow in dense, spatially structured communities where physical interactions matter. We find that under bacteriostatic (growth pausing) antibiotic stress, resistant bacteria can create highly localized protective environments that allow sensitive cells to survive and proliferate. This protection arises not only from antibiotic detoxification, but also from growth–driven mechanical pushing that maintains close cell–cell proximity. As a result, antibiotic tolerance emerges as a collective, spatially dependent property rather than an intrinsic trait of single cells. These findings show that spatial organization, physical forces, and treatment timing can strongly reshape therapy outcomes, with implications for how resistance is understood and managed in microbial communities.

## INTRODUCTION

Bacteria live in complex communities with other prokaryotes, viruses, fungi, and higher eukaryotes. The spatial organization of species is controlled by competition for nutrients and space (1, 2) and cell-cell mechanics (3–6) in a dynamic interplay with chemical interactions. This includes resource exchange (7–9) and responses to environmental stressors, like antibiotics (10–13). This intricate interplay is important for both diversity and ecological stability (14–16).

Increasingly, antibiotic resistance is being explored and understood as a collective property of a multi-strain bacterial community (17, 18). The composition and spatial structure of the community have strong implications for the outcome of antibiotic treatments, leading to diverse and sometimes unexpected dynamics. For example, sensitive bacteria might reduce overall resistance by curtailing the growth of a resistant population (19) or leading to the collapse of the community (20). On the other hand, antibiotic sensitive cells might be protected by their resistant neighbors, who break down the drug and lower the local concentration, an effect called indirect or collective resistance (11, 21).

The outcome for the community and the exact dynamics of indirect resistance depend (among others) on the mechanism of action of the antibiotic and the mechanism by which resistance is conferred. Antibiotics can be either bacteriocidal (i.e., causing lysis) or bacteriostatic (i.e., inhibiting growth). Resistance can be mediated by up-regulation of efflux pumps (22), down-regulation of extracellular influx, mutation of the drug target (23, 24), or enzymatic degradation (25, 26). In the case of enzymatic degradation, the reaction can be extra-cellular (e.g., through secretion of *β*-lactamases to hydrolyze ampicillin or penicillin) or intracellular (e.g., acetylation of chloramphenicol (21)). Secretion of *β*-lactamases can be viewed as ‘altruistic’, reducing the antibiotic pressure across a wider range to enable collective resistance, even in well-mixed cultures (27). In contrast, intracellular deactivation of antibiotics might be considered an ‘egoistic’ strategy of a resistant population. However, sensitive bacteria in their proximity can still benefit from local detoxification to survive otherwise lethal antibiotic concentrations (21). Prior spatial studies of collective resistance have focused on bactericidal antibiotics, where sensitive cells lysed outside of protection zones (10, 11, 13). However, in the presence of bacteriostatic antibiotics, sensitive cells arrest growth and can regrow, if the environment is detoxified. This creates fundamentally different dynamics. Whether and how this difference alters population survival and spatial organization remains unexplored.

Using an *Escherichia coli* surface-attached model system of resistant and sensitive strains and laser-scanning confocal microscopy, we investigated the spatial organization under varying concentrations of the bacteriostatic antibiotic chloramphenicol. We then explored the underlying mechanism by integrating mechanical interactions and antibiotic diffusion and degradation using a lattice model that allows growth, both on the periphery of the colony and deeper in the colony. We demonstrate that growth-driven pushing allows arrested sensitive cells to regrow as a result of local detoxification. In combination, our results show the importance of cells pushing each other towards the periphery of the colony and metabolic interactions (i.e., local antibiotic degradation) for the emerging spatial patterns in a microbial community, providing key insights for the understanding of antibiotic resistance at the community level.

## RESULTS

### Co-culture does not visibly increase fitness of sensitive bacteria

We used two *E. coli* MG1655 strains: a wild type strain (sensitive) and a resistant chloramphenicol acetyltransferase (CAT) producer (resistant), see details in Table 1. We first measured the growth of each strain by optical density, OD_600_, in Luria-Bertani (LB) broth with varying antibiotic concentration, *A*. We found that the maximum growth rate, *λ*, was similar for low concentrations of antibiotics (0-1 µg/ ml) and that the resistant strain was not affected by the antibiotic concentration in the tested range (0-10 µg / ml). In contrast, *λ* of the sensitive strain declined considerably with *A* and could not be distinguished from zero (within uncertainty) beyond 5 µg / ml (Fig. 1A).

**Table 1:**
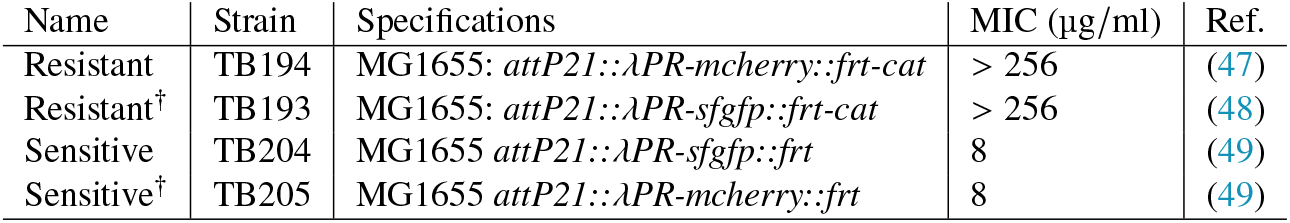
List of *E. coli* K-12 strains used in this study and minimum inhibitory concentration (MIC) of chloramphenicol on agar, as measured using E-test strips. † indicates swapped genes encoding the flourescent proteins. Note that the MIC of the sensitive strain in well-mixed liquid cultures and on agar plates were found to be *∼*5 µg/ml (Fig. 1A) and Supplementary Fig. S2.

**Figure 1:**
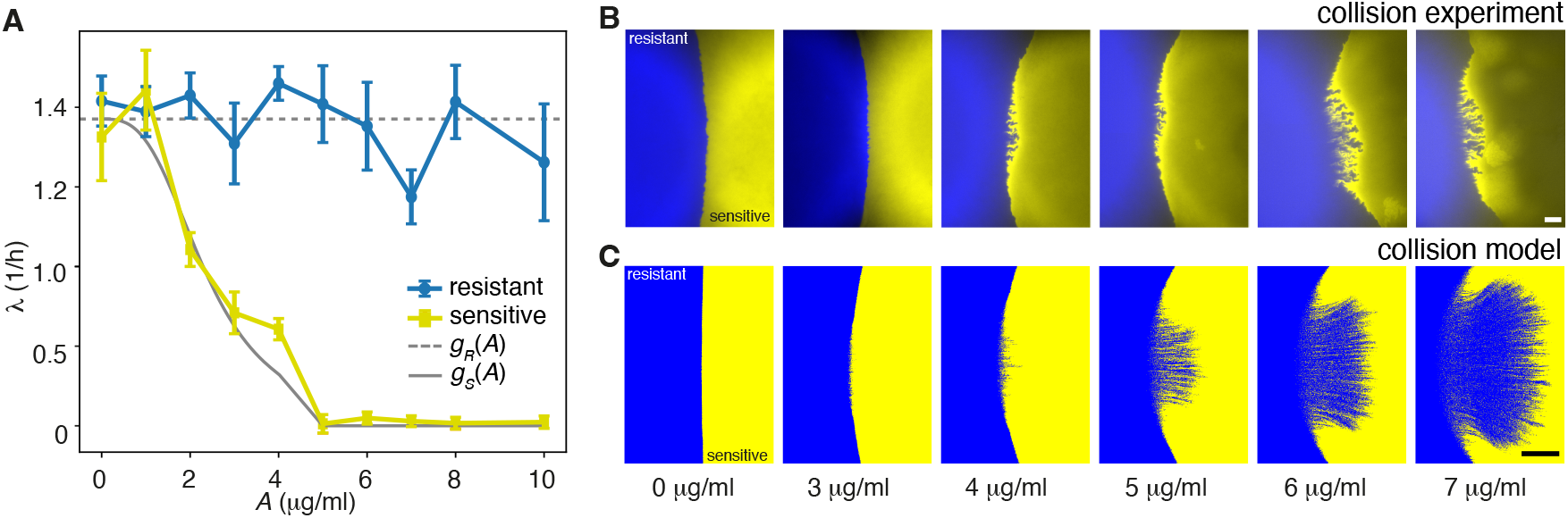
Local detoxification and cell-cell pushing enhance the coexistence of sensitive and resistant bacteria in spatially structured environments. **A:** Maximum growth rates *λ*, of sensitive (yellow) and resistant (blue) in LB liquid media with varying antibiotics concentrations, *A. λ* is obtained from OD_600_ measurements. The gray punctuated line shows the fit to the chloramphenicol independent growth rate of resistant cells *g*_*R*_ *A* = *λ*_*R*_ = 1.93/h. The gray solid line is a fit to a function *g*_*S*_ (*A*) in eq. (4). No significant growth rate differences was found at *A* = 0, so we assumed *g*_*S*_ 0 = *λ*_*S*_ = *λ*_*R*_ = 1.93/h. In addition, no growth of sensitive cells was detected for *A* ≥5 µg /ml and *g*_*S*_ (*A* ≥5 µg /ml) = 0. The fit to the function in eq. (4) for 0 µg /ml< *A* <5 µg /ml gave the half-rate antibiotic concentration: *k* _*A*_ =2.35 µg /ml. **B:** Pseudo-colored fluorescence microscopy images of the colliding front between mono-culture colonies of resistant (blue) and sensitive (yellow) inoculated on agar with varying *A* (see legends). The initial distance between the inoculations was *∼*1 mm (edge to edge) and the colonies were incubated for 24 h. The scale bar corresponds to 200 µm. **C:** Simulated collision experiments using parameters from Table 2. The scale bar corresponds to 200 sites (*∼*200 µm).

To investigate whether the presence of a resistant CAT-producing population has any effect on the growth of the sensitive bacteria, we prepared liquid co-cultures in equal proportions (i.e., the initial ratio of sensitive over resistant bacteria: *χ*_0_ = 1) and different concentrations of chloramphenicol (0-9 µg /ml). Using flow cytometry, we measured the time-dependent fraction of the sensitive population, *ϕ* (*t*), and found that it had a competitive advantage over the resistant population in an antibiotic-free medium. However, this flips completely at high antibiotic concentrations (*A* ≥ 7 µg / ml) (Supplementary Fig. S1).

Having established that there is no considerable advantage for sensitive bacteria to be co-cultured with resistant bacteria in a homogeneous environment, we sat out to investigate antibiotic resistance in a surface-attached bacterial community.

### Local detoxification and cell-cell pushing enhance the coexistence of sensitive and resistant bacteria in spatially structured environments

To test how sensitive bacteria can benefit from direct contact with resistant bacteria in a spatially structured environment, and how the interaction between the two strains influences spatial organization, we inoculated pairs of fluorescently labeled monoclonal colonies with a distance of *∼*1 mm (edge to edge) on LB agar with different concentrations of antibiotics. Within 24 h, resistant and sensitive colonies collided frontally as presented in Fig. 1B. We found that the higher the level of antibiotics, the longer was the collision front, as the populations inter-mingled with one another. To quantify this observation, we measured tortuosity (i.e., deviation from straightness) and intermixing (i.e., crossings with a straight line) of the boundary, confirming that collision fronts became more winding at raised antibiotic levels (Supplementary Fig. S2). We also verified that these results were independent of fluorescence marker gene expression (Supplementary Fig. S3). Furthermore, for higher concentrations (*A* >4 µg / ml), we found a fluorescence gradient in the sensitive population, indicating growth of sensitive cells at the collision front and diminished growth (and fluorescence signal) further away. As chloramphenicol is a bacteriostatic antibiotic and resistance relies on intracellular chloramphenicol-acetylation, our results imply that resistant bacteria locally deplete antibiotics and enable the growth of susceptible bacteria in their vicinity, as was also shown in (21). Also, the self-organized pattern at the community boundary sustain proximity and contact between the two populations.

To understand the nature of this finding, especially the cause of strong intermixing at higher antibiotic concentration, we sat up a computational model (detailed in Methods) to simulate growth and division in a discretized 2D-lattice, where each site represents a bacterium (occupied) or empty space (unoccupied). In contrast to typical Eden-growth type lattice models, where only cells adjacent to empty sites can divide (10, 28), we allowed any cell with unoccupied sites within a radius, *r* _*p*_, to grow and push towards the nearest empty site.

Initially, we placed cells of type *R* (resistant) or *S* (sensitive) within two similar circles with augmented density near the rim, mimicking the so-called ‘coffee stain effect’ of inoculation droplets (29) (detailed in Methods). Then, we let *R* divide with the constant rate, *g*_*R*_, and *S* with the *A*-dependent rate, *g*_*S*_ (*A*), which we obtained by fitting the liquid culture growth experiment (Eq. 4). Antibiotics are assumed to diffuse throughout the lattice with the diffusion constant, *D*, and to be degraded locally at *R*-occupied sites with the rate *γ* · *A*, where *γ* is the intracellular degradation rate. Meanwhile, newly added cells pushed their neighbors towards the closest empty sites, mimicking the intercellular mechanical interactions. Fig. 1C shows the collision fronts after 24 hours of simulated growth, using the default parameter set presented in Table 2. The simulations qualitatively reproduced our experimental results. We found the *A*-degradation rate *γ* to have a strong impact on intermixing (Supplementary Fig. S4) and the default value *γ* = 1.25 × 10^−10^ ml/min was chosen to fit the *A*-dependence of the experimental patterns (Fig. 1B).

**Table 2:** Simulation parameters and scales.

| Parameter | Meaning | Value | Comments |
| --- | --- | --- | --- |
| $x_c$ | Cell lattice site constant | 1 $\mu\text{m}$ | Linear size of an *E. coli* cell |
| $D$ | Antibiotic diffusion coefficient | 200 $\mu\text{m}^2/\text{s}$ | Diffusion constant: 200-1000 $\mu\text{m}^2/\text{s}$ (58–62). |
| $r_p$ | Pushing radius | 50 $\mu\text{m}$ | *E. coli* monolayer expansion (57, 63) |
| $\lambda_R$ | Resistant maximum growth rate | 1.93/h | This work (Fig. 1) |
| $\lambda_S$ | Sensitive maximum growth rate | 1.93/h | This work (Fig. 1) |
| $k_A$ | Half-growth antibiotic concentration | 2.36 $\mu\text{g}/\text{ml}$ | This work (Fig. 1) |
| $x_d$ | Antibiotic lattice site constant | 30 $\mu\text{m}$ | ≪ antibiotic diffusion per generation ($\sqrt{D/\lambda_R}$) |
| $\gamma$ | Resistant antibiotic degradation rate | $1.25 \times 10^{-10}$ ml/min | Fitting parameter (Supplementary Fig. S4) |

To evaluate the importance of cell pushing, we reduced our model to one where only cells next to an empty site can divide (*r* _*p*_ = 1) and without the pushing mechanism, we were unable to reproduce the characteristic pattern of fjords and ridges (Supplementary Fig. S5). Therefore, this alignment between experiments and model intrinsically relies on cell pushing.

Combined with the observed effect of *γ*, we concluded that the intermixed pattern at the collision front is the result of both local detoxification by resistant bacteria and growth-driven mechanical pushing. Specifically in the presence fo antibiotics (*A* > 0), fast-dividing bacteria at the collision front push to squeeze in between sensitive bacteria, while detoxifying the local environment. This allows for enhanced growth of sensitive bacteria, as long as space allows (i.e., unoccupied sites within *r* _*p*_), and reinforces further growth, antibiotics degradation, and pushing.

### Both space to expand and contact to resistant neighbors are necessary for the survival of sensitive populations under antibiotic pressure

To further investigate the role of local detoxification and spatial competition for the survival of sensitive bacteria, we inoculated various ratios of the sensitive and resistant populations (*χ*_0_) at constant combined inoculum density (OD_600_ = 0.3) on LB agar and followed the development of colonies at different antibiotic concentration (1 µg / ml to 6 µg / ml). Fig. 2A shows how for low doses of antibiotics (1 µg / ml column in Fig. 2A), the sensitive strain established sectors (yellow patches) for all *χ*_0_. However, when concentration increased, while maintaining moderate seeding ratios (*χ*_0_ = 2 row in Fig. 2A), sensitive enclaves barely survived at concentrations above 1 µg / ml. In contrast, when sensitive bacteria were much more abundant in the inoculation droplet (*X*_0_ = 10^4^ row in Fig. 2A), they established and persisted to the expanding colony front, even at considerable antibiotic concentrations. These trends demonstrate a strong interaction between antibiotic concentration and initial community composition in determining the survival of a sensitive population.

**Figure 2:**
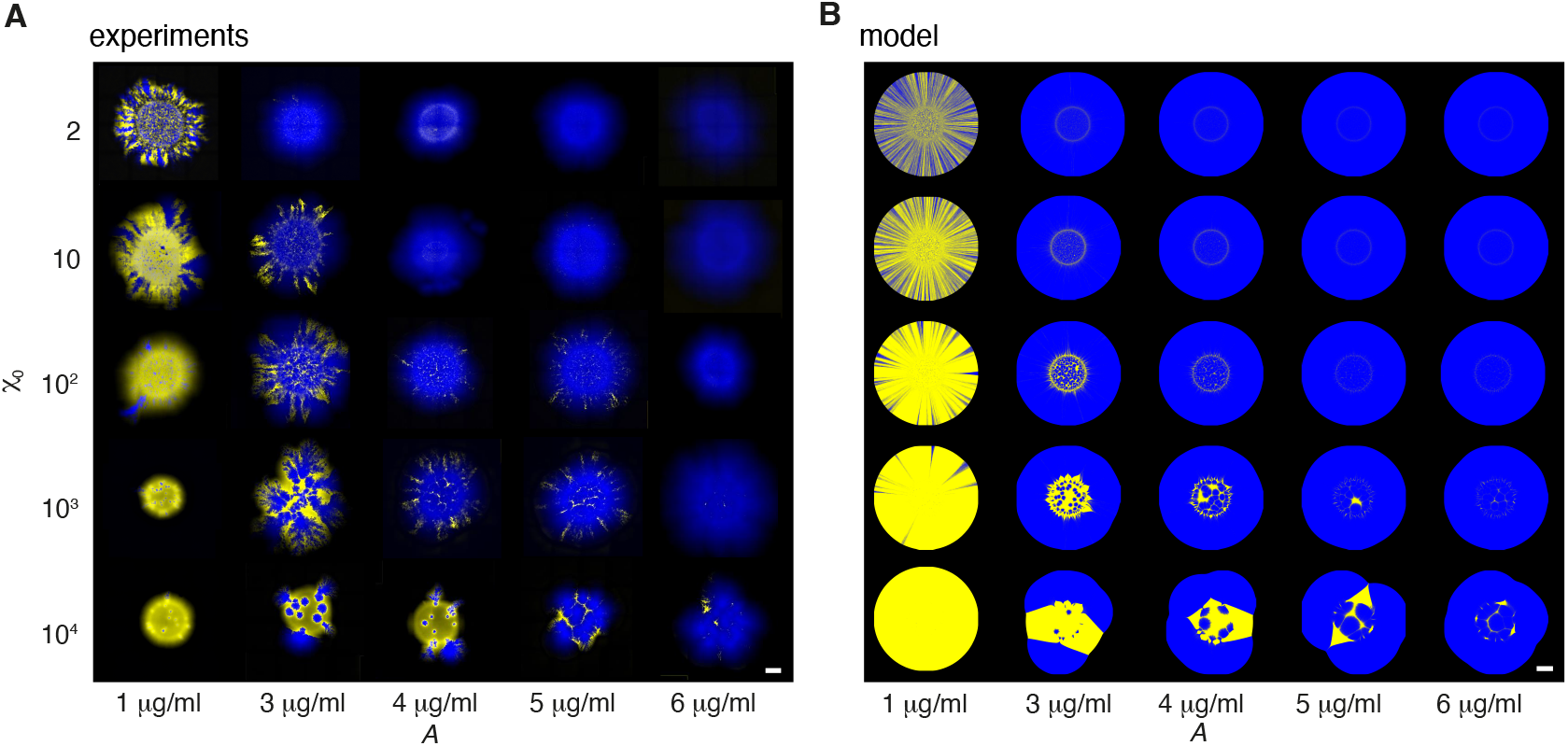
Space to expand and contact to resistant neighbors are necessary for the survival of sensitive populations under antibiotic pressure. **A:** Pseudo-colored confocal laser-scanning microscopy images of surface-attached co-culture colonies of sensitive (yellow) and resistant (blue) at various inoculation ratios *χ*_0_ and inoculation density OD_600_=0.3, on LB-agar with antibiotics *A*. The colonies were incubated for 18 h before imaging. The scale bar corresponds to 1 mm. Note that the minimum-inhibition concentration of sensitive alone is *A* > 4 µg / ml (Supplementary Fig. S2) **B:** Simulated colonies using parameters from (A) found in Table 2. The scale bar corresponds to 1000 sites (*∼*1 mm). The fitness of sensitive bacteria, *w*, in these cases (*χ*_0_, *A*) is given in Supplementary Fig. S6.

The overall stochastic nature of radial range-expansions (3, 30) causes two distinct regions to appear: i) the ‘homeland’, which roughly corresponds to the initial inoculation droplet, and ii) the outermost (i.e., peripheral) region, which is characterized by segregation. A characteristic spatial pattern emerged in the homeland, in which circular domains of resistant cells are surrounded by sensitive cells. This pattern is most pronounced at antibiotic concentrations of 3 µg / ml and 4 µg / ml. Specifically, at a high initial fraction of sensitive (*χ*_0_ = 10^4^ row in Fig. 2A), increasing *A* causes the region of sensitive (yellow) to narrow and the circular resistant (blue) domains to connect, reflecting that rescue of sensitive cells is delayed. As a result, resistant cells expand for longer before sensitive cells can resume growth. A similar merging of circular resistant regions, with narrowing of sensitive regions, is found for fixed *A* when decreasing *χ*_0_ (e.g., 3 µg / ml column in Fig. 2A). In this case, however, the effect arises from reduced initial abundance: when the sensitive fraction is low, more divisions are required before sensitive subpopulations can establish themselves, allowing resistant domains to expand and coalesce before rescue occurs. To test whether these trends are captured by our model, we repeated the experiment *in silico*, matching initial conditions and incubation time and using the parameters identified in the previous section (Fig. 2B). Indeed, we found that the lattice model with mechanical pushing reproduced the emergence of circular resistant regions. At high concentrations of antibiotics (*A* ≥ 5 µg / ml), only resistant cells grew at early times. Their proliferation mechanically displaced sensitive cells outward. As the environment became sufficiently detoxified, sensitive cells resumed growth. This sequence naturally generated resistant cores surrounded by sensitive cells and highlights the importance of mechanical interactions in spatial organization. The establishment of sensitive cells in the homeland, therefore, depended both on the time to rescue by detoxification (controlled by *A*) and on the initial abundance (controlled by *X*_0_). These factors directly affect the probability of surviving long enough to reach the periphery and continue growing with colony expansion.

To quantify the impact of this spatial bottleneck, we calculated the relative fitness, *w*, of the sensitive population with respect to the resistant (Eq. 2). We confirmed that fitness decreased with increasing antibiotic concentration, but that this decrease was substantially mitigated by a higher seeding ratio *χ*_0_ (Supplementary Figure S7), increasing the probability of rescue. Our model also showed qualitatively consistent behavior: (i) for *A* =1 µg / ml, sensitive sectors established and expanded; (ii) for *A* = 3 −4 µg / ml, sufficiently large *χ*_0_ led to some sensitive sectors persisting to the expanding periphery; (iii) at higher *A* or lower *χ*_0_, circular resistant regions merged, leaving only narrow sensitive regions, and sensitive sectors at the periphery eventually closed off. Moreover, relative fitness, *w*, followed these same general trends (Supplementary Fig. S7).

These observations indicate that local degradation alone is not sufficient to guaranty long-term survival. The resistant population can detoxify enough to revive the sensitive bacteria. However, for the sensitive population to be established and sustained, it must reach (and stay on) the colony’s expanding periphery, where unoccupied space and nutrients are available.

### Proximity favors the rescue of sensitive cells

We hypothesized that *χ*_0_-dependence reflects the need for sensitive bacteria to stay close to resistant cells to benefit from their local clearing of antibiotics. To verify this, we repeated the experiment of Fig. 2A with a higher inoculum density (OD_600_ = 3), while keeping the initial ratio constant (*χ*_0_ = 10^4^). We found that lower seeding density (OD_600_=0.3) is advantageous for the sensitive population at ambient antibiotic concentrations (*A* ≤ 4 µg / ml), as it ensures the required space to establish expanding regions. However, this trend shifts at higher concentrations (*A* =5 µg / ml), where sensitive bacteria rely on detoxification by neighboring resistant cells in order to proliferate. In this regime, higher seeding density gave more visible growth, likely because reduced cell-cell distances increased the likelihood that sensitive cells benefit from the local *A*-degradation. This shift is quantified by calculating the relative fitness *w* of the sensitive bacteria in the periphery of the colony (Fig. 3B).

**Figure 3:**
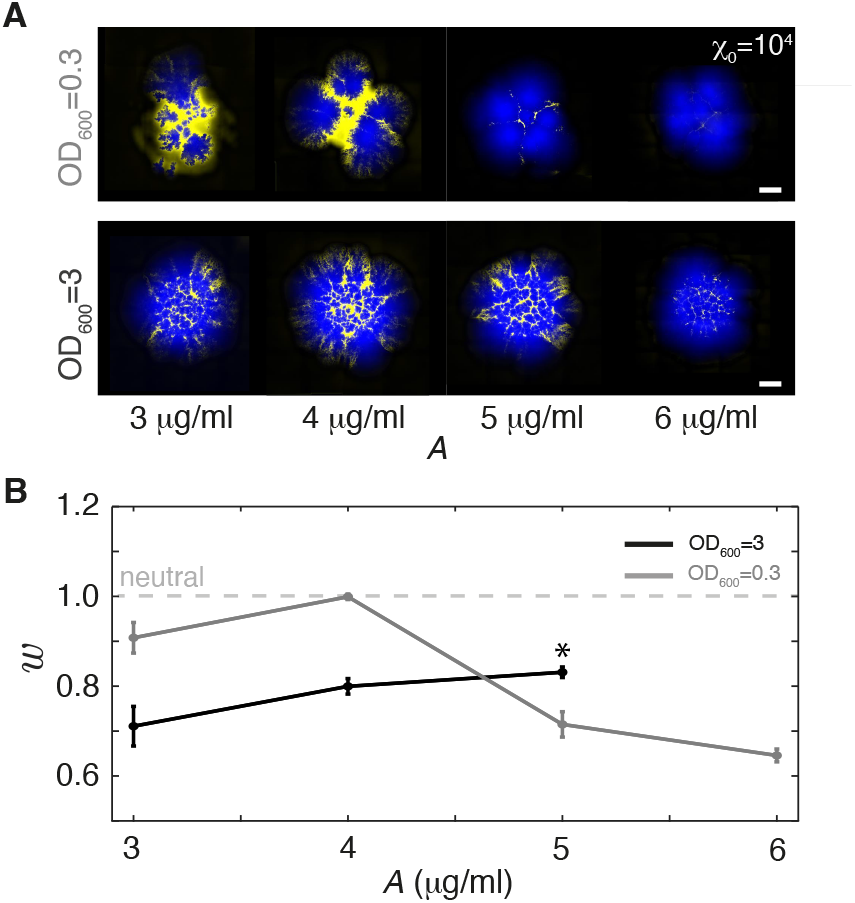
Proximity favors the rescue of sensitive cells. **A:** Pseudo-colored confocal laser-scanning microscopy images of surface-attached co-culture colonies of sensitive (yellow) and resistant (blue) at inoculum ratio *X*_0_ = 10^4^ and densities of either OD_600_=0.3 (similar to Fig. 2) or OD_600_=3 on LB-agar plates with different antibiotic levels, *A*, incubated for 18 h. The scale bars correspond to 1 mm. **B:** The average fitness of the sensitive strain relative to resistant, *w* (as defined in Eq. 2), at various inoculum densities, OD_600_, see legends. * signifies the last *A*-level at which the sensitive was detected (i.e., *w* diverges at *A* = 6 µg/ml). The horizontal dashed line corresponds to equal fitness (*w* = 1) and the error bars are SEM (*N* = 2 for all entries).

At ambient antibiotic concentrations (*A* ≤ 4 µg / ml), low seeding density (OD600 =0.3) ensures that sensitive patches can grow large enough to form sectors capable of reaching the periphery. However at *A* =5 µg / ml, survival is considerably enhanced for higher seeding density (OD_600_ =3). At *A* =6 µg / ml, the *w*-trend was unclear (Supplementary Fig. S8), because the overall probability of the sensitive population reaching the periphery is low, despite pronounced homeland growth at OD_600_ = 3 (Fig. 3A, Supplementary Fig. S8A). Still, these results support the interpretation that detoxification is local and that sensitive cells must be in the immediate vicinity of resistant cells to be rescued (i.e., resume growth) under high antibiotic pressure.

### Administration time determines proliferation of the sensitive population

The importance of the inoculum density for the survival of sensitive cells at high antibiotic concentrations indicated the need for spatial proximity to resistant neighbors. We next asked how antibiotic stress reshapes spatial structure once sectors have already formed at the expanding frontier, independent of early-establishment effects.

To address this question, we mixed sensitive and resistant bacteria (*χ*_0_ = 2) and allowed colonies to expand in the absence of antibiotics. After 6 h, indicated by the white dashed line in Fig. 4A, well-defined wide sectors had established and antibiotics were added (3 µg / ml or 5 µg / ml). Then sensitive sectors gradually narrowed in and closed off, while the colony continued radial expansion. The merging sector boundaries (outside white dashed line) were relatively smooth, compared to the meandering sector boundaries before the addition of antibiotics (within white dashed line), reflecting the deterministic drift of the sector boundary towards loss of sensitive sectors. This behavior demonstrated that even wide sensitive sectors are unstable under antibiotic stress. The progressive lateral encroachment of resistant sectors indicated that detoxification cannot sustain sensitive cells across large distances. Instead, sensitive bacteria far from the sector boundaries will be too distant from resistant neighbors to benefit from their detoxification, leading to systematic sector narrowing. The requirement of proximity for rescue creates a trade-off for sensitive cells at the colony front: If the sensitive sector is too wide, bacteria deep in the sector are growth-inhibited by high antibiotic, causing the sector to narrow rapidly. Conversely, when a sector narrows, a fraction of sensitive bacteria can make close contacts and benefit from local detoxification, potentially decelerating the thinning of the sector.

**Figure 4:**
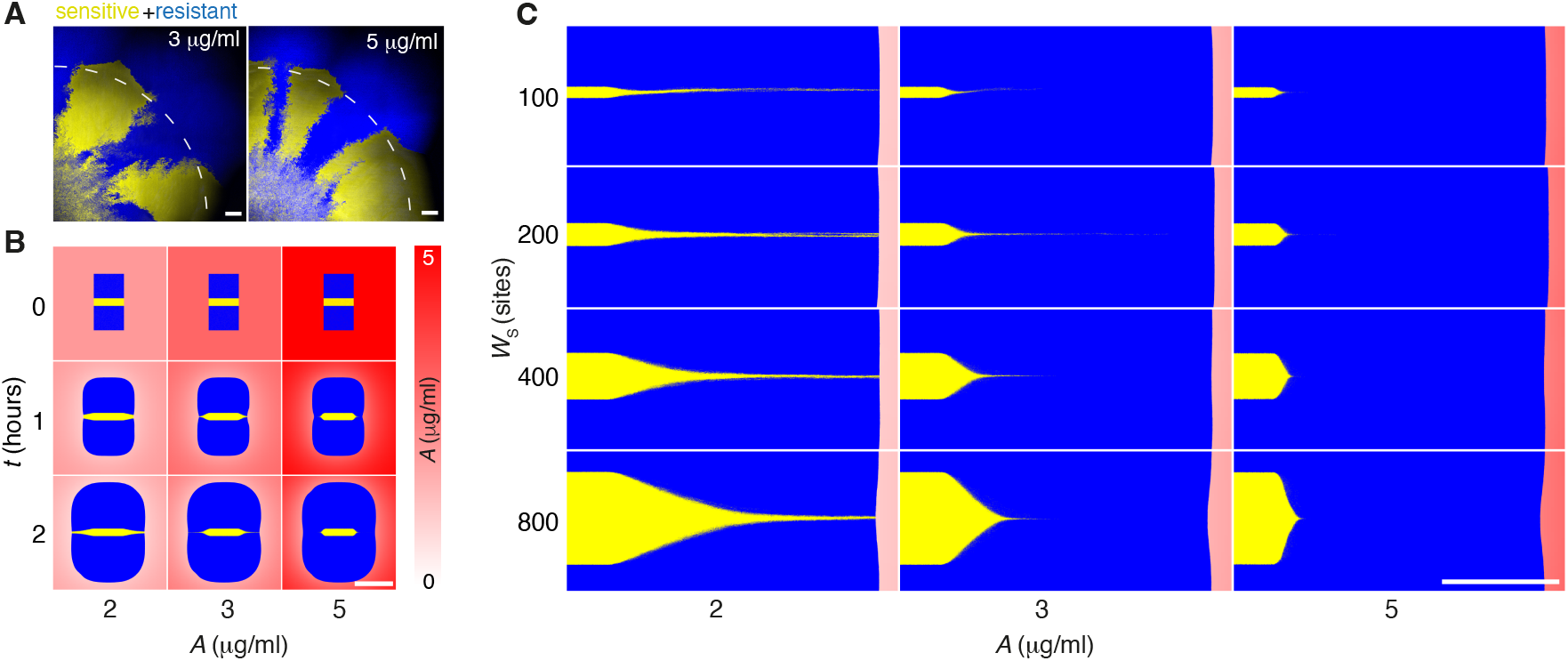
Late chloramphenicol administration changes fitness. **A:** Pseudo-colored confocal laser-scanning microscopy images of surface-attached co-culture colonies of sensitive (yellow) and resistant (blue) inoculated at *X*_0_ = 2 and a density of OD_600_ = 0.3 on LB agar with an antibiotic concentration of either *A* =3 µg / ml or *A* =5 µg / ml. Samples were incubated for 6 h for sectors to establish before antibiotics were added as droplets in a circle around the periphery and then incubated for another 18 h. The scale bars correspond to 200 µm. **B:** Simulation time courses of rectangular colony inocula of sensitive sectors (yellow) with the width of *W*_*S*_ = 200 sites sandwiched by resistant (blue) sectors. The background color indicates the antibiotic concentration, *A*. **C:** Magnification of simulated colonies at 8 h for different *W*_*s*_ and *A*. Scale bars in B and C corresponds to 1000 sites (*∼*1 mm)

To test this, we simulated colonies starting from a rectangular region (800 × 1500 sites) with 90% occupancy. The inoculum (*t* = 0) was composed of a central sector of type *S* of width *W*_*S*_, flanked by type *R* in the uniform field of concentration *A*. Fig. 4B shows how colony expansion and the narrowing of the sensitive sectors over time depended on initial concentrations: *A* = 2, 3, and 5 µg / ml, which is degraded over time. Fig. 4C shows magnified views after 8 h for different *A* and *W*_*S*_ = 100, 200, 400, and 800 sites. For wider sections of sensitive cells, the boundary between resistant and sensitive cells were angled, indicating ongoing closure, as the resistant population progressively overtake the sensitive. As the initial width decreased, the angle became shallower, reflecting slower closure rates. At *A* =2 µg / ml, the sensitive sectors reached a (nearly) constant width, indicating that closure has effectively stopped. In these cases, sensitive bacteria benefitted from continuous detoxification, allowing them to persist to the colony front. However, only 15± 2 sensitive cells remained at the colony edge after 8 h (mean ±SEM; left and right edges counted separately across 8 independent simulations of different *W*_*S*_, *N* = 16). While the closure rate is slow, the few surviving cells suggest that stochastic demographic fluctuations will eliminate the sensitive population over longer timescales.

## DISCUSSION

In summary, our results show that indirect antibiotic resistance in structured bacterial communities is an interplay between spatial organization, local detoxification, and cell mechanics. In our *E. coli* system, exposed to chloramphenicol, resistant bacteria create favorable micro-environments that allow sensitive neighbors to resume growth, while colony expansion and pushing maintain the close cell-cell proximity required for this rescue.

Our work highlights a key distinction between bacteriostatic and bactericidal antibiotics. Because chloramphenicol suppresses growth, rather than causing lysis, sensitive cells can recover when concentrations are reduced. This leads to complex dynamics, which are consistent with reports of non-trivial dynamics in well-mixed liquid cultures (20, 21, 27) and in surface-attached communities, where the feedback between local antibiotic inhibition and mechanical displacement can either promote one population or generate persistent coexistence (12, 31–33). This explains why intermixing, homeland structure, and sector closure are not merely visual features of colony growth, but direct determinants of community-level fitness (Fig. 2). The intermixing index (Fig. 1) is, thus, a key quantitative measure of coexistence: higher intermixing enhances the protection of sensitive bacteria. A similar connection between intermixing and inter-dependence has been reported for communities with metabolite exchange (i.e., cross-feeding), both in bacterial (9, 34, 35) and yeast systems (36).

Our results also emphasize the importance of cell-cell pushing, which was evident when fine channels of resistant bacteria squeezed into regions initially populated by the sensitive population (Fig. 1), when circular resistant regions formed narrow sensitive channels (Fig. 2, e.g., *A* = 3 µg / ml, *χ*_0_ = 10^4^), or when thin sectors of the sensitive population persisted to the expanding front (Fig. 3). In conclusion, mechanical pushing leads to very local protection zones, reinforcing proximity between sensitive and resistant strains. This is in contrast with long-range rescue scenarios arising from antibiotic degradation by *β*-lactamases (10, 11, 13).

Despite a high degree of qualitative correspondence, our model overestimated the radial expansion of surface-attached communities (Fig. 2). This discrepancy arose mainly from dimensionality, as on a two-dimensional lattice, growth always caused an area increase. In contrast, in the experimental system, bacteria built new cell-layers (i.e., grew into the third dimension), slowing radial expansion. Furthermore, our model did not reproduce the strong meandering of sector boundaries (Fig. 1 and 2), caused by growth-driven pushing of rod-shaped cells (dividing along their long axis) and generating interfacial instabilities (37–39). Indeed, this limitation could be addressed by individual-based modeling of cell shape and mechanical interactions (6, 12). Instead, we chose a minimal model and kept computation simple by adding growth-driven pushing to a lattice model. Despite the simplicity, our model captured the emergence of indirect resistance and the resulting growth dynamics of the surface-attached community and showed contingency in both initial community composition and antibiotic exposure.

From optical density measurements, we found equal growth rates for sensitive and resistant strains in an environment without antibiotics (Fig. 1A), which we used as input to the model. However, the minor curvature observed on the collision front without antibiotics (Fig. 1B, *A* = 0) and our flow cytometry data (Fig S1, *A* = 0) indicated a slight resistance cost. Furthermore, this cost seemed to disfavor the resistant strain even at low antibiotic levels, as indicated by the flip in curvature between *A* ≤ 3 µg/ml and *A* ≥ 4 µg/ml(Fig. 1B).

Engineered communities with mutual or unidirectional strain dependencies have been found to converge to similar population ratios over time, despite different initial conditions (36). For example, in well-mixed liquid cultures, the sensitive-resistant ratio stabilizes if sensitive cells grow faster at low antibiotic levels; while high fractions of resistant cells enhance detoxification, excess sensitive cells diminish this effect (27), resulting in a final ratio that balances the trade-off. Similar trade-offs may stabilize sensitive survival at the expanding front. As, faster growth of sensitive cells, enabled by local detoxification, causes a loss of proximity to resistant cells, detoxification is slowed down and further growth of sensitive domains inhibited. The requirement for proximity to resistant cells underscores the importance of timing of antibiotic treatment, as the typical inter-strain distance at the growing front expands with time (40).

In complex microbial communities, member species can produce antimicrobial compounds that target others (41), altering system dynamics and underscoring the importance of understanding the impact of antibiotics on spatial organization, beyond therapeutic antibiotic applications (42). Also, the organization has consequences for a variety of processes, such as horizontal gene transfer (43, 44) and biofilm formation (45, 46). Together, the coupling between local detoxification and spatial diversity enhances indirect resistance. Thus, a key outcome of our study is establishing a causal pathway linking local antibiotic degradation, mechanical interactions, spatial organization, and colony-level tolerance.

## MATERIALS AND METHODS

### Cell culture

#### Bacterial strains

We used the set of *E*.*coli* MG1655-derived strains listed in Table 1 with constitutive expression of either red fluorescence (mCherry) or superfolder green fluorescent protein (sfGFP), where sensitive (sensitive^†^) denotes the wild-type strains sensitive to chloramphenicol and resistant (resistant^†^) the strains with *cat* gene encoding chloramphenicol acetyltransferase (CAT) that inactivates chloramphenicol by acetylation. Strains are kind gifts from the laboratory of Olga T. Schubert at Eawag - Swiss Federal Institute of Aquatic Science and Technology, Zürich.

#### Culture medium

We used rich Lysogenic broth (LB) composed of 10 g l Gibco™ Bacto™ tryptone (16279751, Fisher Scientific), 5 g l Gibco™ Bacto™ yeast extract (16279781, Fisher Scientific) and 5 g l NaCl (≥ 99%, S9888, Sigma-Aldrich) dissolved in MilliQ water, supplemented with 1.5 % (w/v) Bacto™ agar when working with plates.

### Competition in liquid co-cultures

#### Flow cytometry

Starting from overnight cultures, we prepared a set of mono-cultures (resistant and sensitive), as well as a co-culture mixed in equal ratios (*χ*_0_ = 1) based on OD_600_ measurements. These three cultures were each diluted 100 × in LB and incubated (37 °C) with various concentrations of chloramphenicol. Cell counts and fluorescent signals (50000 events per sample) were measured instantaneously after dilution and again after 6 h of incubation, using a BD FACSJazz Cell Sorter (BD Biosciences, cat. no. 655490). Data were processed using the Flowkit package (50) with a *hyper-log* transformation, and clustered with the *hdbscan* algorithm (51). After clustering, we extracted the counts of the two main clusters (mCherry and sfGFP) and the counts, which were not included in a cluster, were < 2% in all cases. Finally, we defined the time-dependent frequency of a population, *ϕ*(6 h), as the ratio of cell counts in one cluster over the sum of both clusters after 6 h.

#### Relative fitness

The fitness of a strain under given environmental conditions can be quantified as a selection coefficient at which sensitive bacteria are eliminated by resistant bacteria at each division (52):

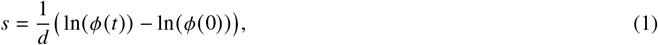

where *ϕ*(0) is the initial frequency at *t* = 0 of the sensitive population and *d* is a scaling factor, corresponding to the expected divisions over the full growth cycle (i.e., log_2_ *d* of the fold-change after *t*). From *s* the relative fitness can be found:

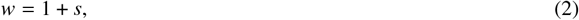

which is a measure of the strength, with which the sensitive strain is enhanced (diminished) in co-culture with the resistant if *w* > 1 (*w* < 1).

#### Growth rate measurements

From single colonies, we inoculated four individual precultures of resistant (TB194) and sensitive (TB204) in LB medium. After overnight incubation at 37 °C, we measured OD_600_ and normalized all precultures to OD_600_=0.3 using LB medium. 5 µl of the normalized precultures were used to inoculate growth cultures in a final volume of 150 µl in LB medium containing the appropriate concentration of chloramphenicol. This is equivalent to a total dilution of approximately 1:500 from the overnight cultures. We prepared four replicate cultures of each of the two strains for various concentrations of chloramphenicol (0 − 10 µg / ml). OD_600_ measurements were recorded every 5 min using a FLUOstar Omega microplate reader (BMG Labtech, Germany) under continuous shaking over a period of 24 h.

#### Minimum inhibitory concentration

We found the minimum inhibitory concentration (MIC) by spreading 200 µl bacterial cultures of OD_600_ = 0.2 (IMPLEN NanoPhotometer C40) on LB agar plates (1.5%) and carefully placing an E-test strip (Biomerieux, France). The plates were incubated for 20 hours at 37°C before estimating (by visual inspection) the chloramphenicol concentration, at which growth was inhibited (Table 1). We measured an alternative minimum-inhibition concentration as the growth detection limit at > 4 µg/ml (Supplementary Fig. S2).

### Collision fronts between mono-strain surface-attached colonies

Overnight cultures (resistant+sensitive) were each diluted in LB to an OD_600_ = 0.3. Droplets of 1 µl of each strain (OD_600_ = 0.3) were inoculated, with approximately 1 mm distance, on 1.5% LB agar containing the specified chloramphenicol concentrations. Six pairs of droplets were inoculated per dish and incubated at 37 °C for 24 h. Colliding colonies were imaged using an inverted fluorescent microscope (Nikon ECLIPSE Ti microscope) paired with a 4 × air objective (Nikon, Plan Fluor, 4 ×/0.13, ∞ /1.2 WD 16.5). The fluorescent signals were recorded sequentially, using a Hg lamp and either FITC (480/40 nm) or Texas Red (560/40 nm) cubes in combination with a camera (Andor Neo, Oxford Instruments). Pre-processing and thresholding was done with Fiji ImageJ (53).

#### Tortuosity and Intermixing index

A common (all images, all samples) region-of-interest was chosen and the GFP-fluorescence channel (corresponding to sensitive) was thresholded to produce a salt-and-pepper image of the footprint of the sensitive strain. The collision front was found using an edge detection procedure and we defined the tortuosity, *T*, as the ratio between the contour length of the edge and the length of the line connecting its end-points. The same salt-and-pepper image was line-scanned (along the direction parallel to the edge) measuring the number of intersections with the collision boundary. The maximum number of intersections per length is the intermixing index, *I*.

### Range expansion in surface-attached co-culture colonies

Overnight cultures were mixed in various ratios of sensitive over resistant bacteria, *χ*_0_ *∈* {2, 10, 10^2^, 10^3^, 10^4^}, to a final (and combined) OD_600_ *∈* {0.3, 3}. The diluted cell culture was inoculated in 1 µl droplets on LB agar plates (1.5%) with various concentrations of chloramphenicol (0− 6 µg / ml), and incubated for 18 h-48 h (37 °C). After incubation, we imaged the colonies using a confocal laser-scanning microscope (DMI6000 CS SP5, Leica, Wetzlar, Germany). We used the Ar (488 nm) and HeNe (543 nm) laser lines sequentially to excite sfGFP and mCherry, respectively, using a 5× air objective (Nplan5× 0.12PHO, Leica). We made a maximum intensity pixel projection of each stack (corresponding to each fluorescence channel) to get a flattened (*x, y*) -image before stitching. The resulting image had the pixel size (*x, y*) = (6.07, 6.07) µm. In the cases where the colonies exceeded the field of view (3.1 mm × 3.1 mm), we stitched four images together using the Pairwise Stitching plugin in Fiji ImageJ (54).

#### Relative fitness in surface-attached colonies

Colonies exhibited clearly distinguishable regions occupied by sensitive and resistant populations. To quantify the relative fitness at the end of the incubation period, we performed the following image-analysis procedure. The projected image of the sensitive strain (GFP channel) was thresholded to generate a binary (i.e., salt-and-pepper) image identifying pixels occupied by sensitive cells. In parallel, the projected images from both fluorescence channels (GFP and mCherry) were summed to obtain the footprint of the entire colony, from which a one-pixel-wide binary map of the outer rim of the colony was extracted. The rim mask was then overlaid with the binary sensitive image by pixel-wise multiplication. Rim pixels overlapping with the sensitive mask were classified as sensitive, while the remaining rim pixels were classified as resistant. This yielded coarse-grained estimates of the number of bacteria belonging to the sensitive and resistant populations, denoted *N*_S_ and *N*_R_, respectively. Assuming that the outer rim represents the most recent cell growth, the frequency of sensitive cells after the total incubation time (18 h) was calculated as

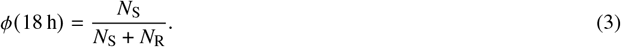

Using these frequencies, we calculated the selection coefficient (Eq. 1) at the periphery for each division. However, here we used the number of divisions, *d* = 13 (*∼ λ*_*S*_/18 h), as scaling factor to get *w* (Eq. 2). We took a similar approach to analyze the modeling data: Locating the outer rim and calculate *N*_S_ and *N*_R_. To avoid divergence *w* was only defined for *ϕ*(18 h) > 0.

### Range-expansion and late chloramphenicol administration in surface-attached colonies

Overnight cultures were mixed in equal proportions (*χ*_0_ = 2) based on OD_600_ measurements to a final (and combined) OD_600_ = 0.3. The diluted cell culture was inoculated in 0.5 µl droplets on LB agar plates (1.5%) in the absence of chloramphenicol and incubated for 6 h (37 °C). Hereafter, chloramphenicol was added following the protocol of Ref. (55) by inoculating 4 droplets with 10 mg ml chloramphenicol on a circle of radius 50 mm around the center of the colony for a total concentration of 3 µg / ml or 5 µg / ml. Then, samples were incubated for another 18 h (37 °C) before imaging using the settings detailed above.

### Lattice model to simulate indirect resistance on a two-dimensional surface

We further explored the system using a stochastic square-lattice model. The cells resided in a unit step size lattice (1µm), with each lattice site being 0 (free) or 1 (occupied). The model represents cell division, expansion of the colony, antibiotic diffusion and degradation, and the effect of antibiotic on the cell division as follows.

#### Cell types and growth

We introduced two different cell types, the sensitive *S* and resistant *R*. While *R* grew at a constant rate *g*_*R*_ (*A*) = *λ*_*R*_, *S* was inhibited by the local concentration of antibiotics, *A*, in units of µg/ml (56):

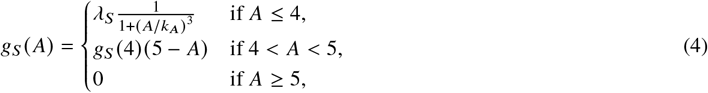

where *g*_*S*_ 0 = *λ*_*S*_ is the maximal growth rate (no antibiotics) and *k* _*A*_ the half-rate constant, determined by least-squared fitting of sensitive growth rate when varying *A* (Fig. 1A). Both cell types progressed toward division using a 10-step growth process, where cells divided after completing the 10 steps. In each time step, a cell can progress one step toward division with probabilities *p*_*R*_ = 10(*g*_*R*_ (*A*)/ln 2)*dt* and *p*_*S*_ = 10(*g*_*S*_ (*A*)/ln 2)*dt* for a resistant cell and a sensitive cell, respectively. The 10-step process ensured the inter-division time distribution peaked around the average inter-division time with coefficient of variation 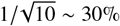, in contrast to one-step division where the inter-division time distributes exponentially.

#### Diffusion and degradation of antibiotics

The diffusion of antibiotics through the agar was modeled according to the two-dimensional diffusion equation

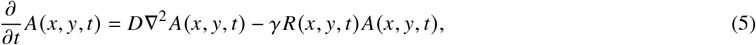

where *D* is a constant diffusion coefficient *D, γ* is the antibiotic degradation rate, and the last term is the local degradation of *A* at *R*-occupied lattice sites; *x* and *y* denote the location on the square lattice and *t* the time. This was based on the assumption that *A*-degradation is limited by collision frequency between the *R* cell and diffusing antibiotic molecules, rather than enzyme-antibiotic binding kinetics or diffusion of antibiotic molecules across the cell membrane. We assumed that the antibiotic field only changed in the horizontal plane (along the agar surface) and was constant in the vertical direction (in depth). As the depth of the system was assumed (and estimated) to be 0.5 cm, a cell lattice site of 1*μ*m^2^ area had a volume of 5× 10^3^ µm^3^ = 5× 10^−9^ ml.

The spatial size of the antibiotic lattice was set to 30 cell lattice sites. As antibiotic diffusion in agar is fast on the cell length and time scale, the antibiotic gradient will be small, making a coarser lattice an efficient choice without losing necessary resolution to follow the system’s dynamic. The diffusion equation was solved using the implicit Euler method with a 5-point stencil and optimized using an incomplete LU precondition.

#### Cell division and pushing

When cells divided, the nearest empty lattice sites within pushing radius, *r*_*p*_, were identified. If no unoccupied site was found, the cell did not divide. This effectively takes into account cell division limitations (i.e., lack of nutrients and space) when a cell is far from the colony edge. It also reflects the limit in horizontal expansion due to friction, though we did not explicitly simulate the formation of multi-layers (57).

If a free lattice site was found within *r*_*p*_, a sequence of moves in *x* and *y* was used to determine a path between the dividing cell and the free lattice site based on the probability

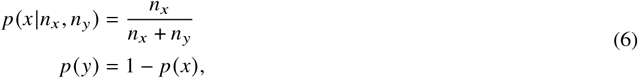

where *n*_*x*_, *n*_*y*_ are the number of lattice sites separating the two sites in the respective directions. *n*_*x*_, *n*_*y*_ are adjusted accordingly after each choice, making the sequence generated analog to sampling without replacement.

Once the pushing path was determined, the cells were moved accordingly, after which a copy of the dividing cell was placed on the newly freed lattice site. This very simple rule efficiently represents the repulsive mechanical interaction between cells, which allowed us to explore large colonies using cell counts identical to the experiments.

#### Initial condition

To simulate the collision experiment (Fig. 1, Supplementary Fig. S4) and the range expansion of co-cultures (Fig. 2, Supplementary Fig. S6), er initially deposited the cells, as to mimic inoculation droplets with augmented density near the rim (i.e., coffee stain effect (29)). 20% of the occupied sites (*R* and/or *S*) were uniformly distributed within a circle of 1000 sites (*∼*1 mm). The remaining 80% were arranged in a band between circles of radius 1000 sites and 1100 (*∼*1.1 mm). The total number of cells in the initial condition was calculated by assuming that a culture of bacteria with OD_600_ = 1 contained 2.5× 10^8^ cells/ml.

To simulate the antibiotic addition to an established sector (Fig. 1), a 1500 × 800 sites rectangular region was filled by cells with 90% site occupancy. The central region of width *W*_*S*_ contained sensitive cells, and the rest was filled by resistant cells. An uniform antibiotic concentration *A* across the system was imposed at *t* = 0.

#### Parallelization and order of operations

The pushing radius, *r*_*P*_, corresponds to an interaction distance for a given cell within a time step. Thus, different regions of the lattice can be time-stepped simultaneously, as long as they are separated by a distance > 2*r*_*P*_. For each time step, the cell lattice is separated into a number of these regions. Then, for each of these regions, the cell growth, division, pushing, and antibiotic degradation are computed, after which the diffusion equation is solved for the entire grid.

## AUTHOR CONTRIBUTIONS

MC, NM, and LJ conceived the original idea. AKE, MC did liquid culture experiments. MC and AT did surface-attached colony experiments. BHT and NM developed the model. BHT developed the algorithm and performed simulation and data visualization. SLS contributed to the conceptual development of the experimental approach and early experimental development. NM and LJ supervised the project.

## DATA AVAILABILITY

The simulation code is available at https://github.com/namikomit/CellColonyLatticeModel.

## FUNDING

This work was supported by the Danish National Research Foundation grant no. DFF 0165-00032B and grant no. DFF 0165-00103B (LJ), the Novo Nordisk Foundation NERD grant no. NNF21OC0068775 (NM) and Synergy grant no. NNF23OC0086712 (LJ).

## ACKNOWLEDGMENTS

The authors thank Adrien Sarlet for his help with wide-field microscopy.

## Notes

### Competing Interest Statement

The authors have declared no competing interest.

