## Supplementary Information for "Pushing for survival: Spatial intermixing and indirect resistance enable collective growth"

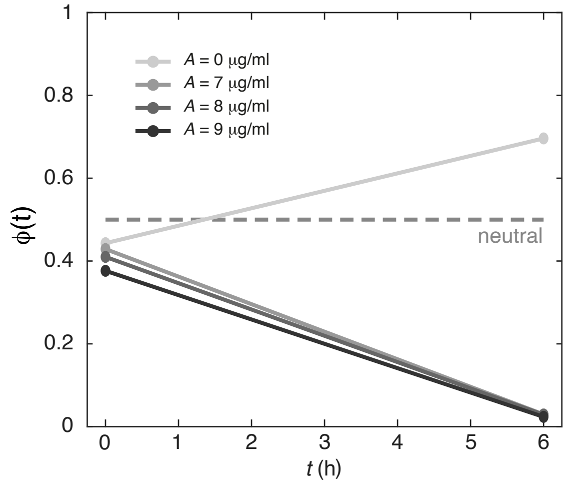


**Figure S1**. Growth versus antibiotics concentration. Competitions between equal ratios ($\chi_0 = 1$, based $OD_{600}$ of resistant and sensitive in liquid LB medium under constant shaking. The frequencies of the sensitive in the co-culture at time $t$: $\phi(t)$ at various Cm concentrations, see legends. The data are obtained using flow-cytometry and the dotted line corresponds to the neutral case (i.e., $\phi(t) = 0.5$).

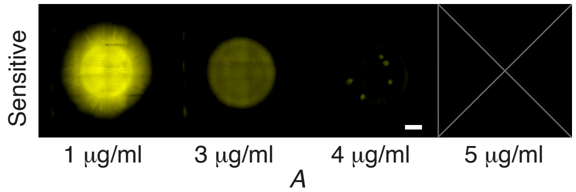


**Figure S2**. **Growth of sensitive bacteria.** Pseudo-colored confocal laser-scanning images of surface-attached sensitive (yellow) and $OD_{600}$=0.3 on LB-agar with different Cm levels. The colonies were incubated for 18 h before imaging. No growth was detectable for [Cm]> 4 μg/ml. The scale bar corresponds to 1 mm.

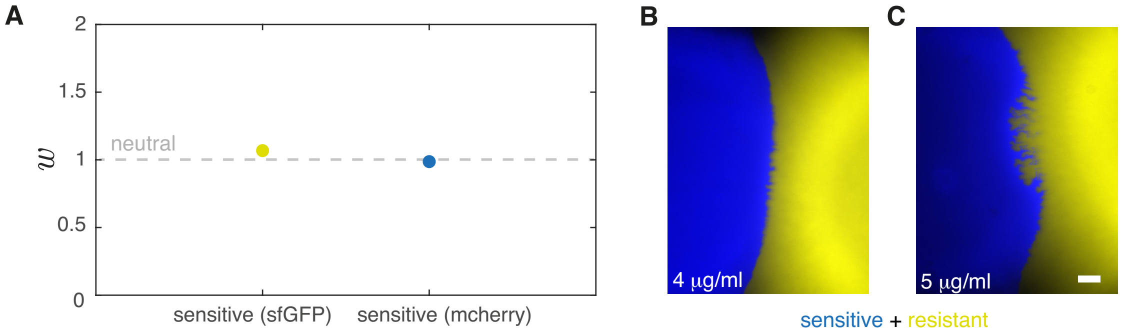


**Figure S3**. **Fluorescence color does not change competitive behavior. A:** Competitions between resistant and sensitive in liquid LB medium (no Cm) under constant shaking and for swapped colors. The average fitness of the sensitive (sensitive$^\dagger$) in respect to the resistant (resistant$^\dagger$)strain, $w$, after 6 h. The data are obtained using flow-cytometry ($\chi_0$ = 1) for the strain set used through-out the study, sensitive (sfGFP), and the set with swapped colors, resistant$^\dagger$ (mcherry). $w$ = 1 corresponds to equal fitness. **B-C:** Pseudo-colored fluorescence microscopy images of the colliding front equivalent to Fig. 1A in the main text with swapped colors. Colonies of sensitive$^\dagger$ (blue) and resistant$^\dagger$ (yellow) were inoculated (~1 mm apart) on an agar surface with either [Cm] = 4 μg/ml (B) or [Cm] = 5 μg/ml (C). The colonies were incubated for 24 h. The scale bar corresponds to 200 μm.

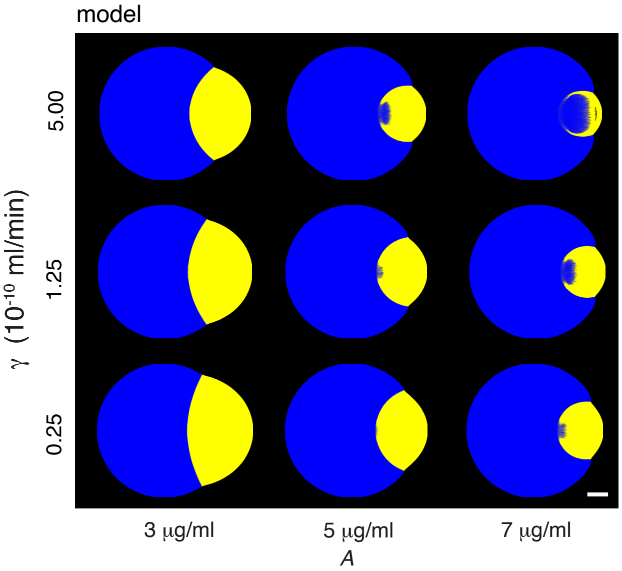


**Figure S4**. **Antibiotic degradation, $\gamma$, versus antibiotic concentration, $A$.** The scale bar corresponds to 1 mm.

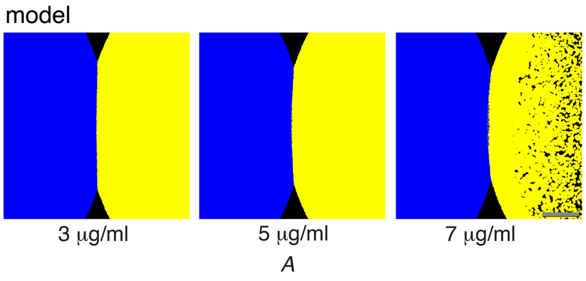


**Figure S5. Local detoxification without pushing decrease coexistence of sensitive and resistant bacteria.** Antibiotic concentration, $A$. Scale bar corresponds to 1 mm.

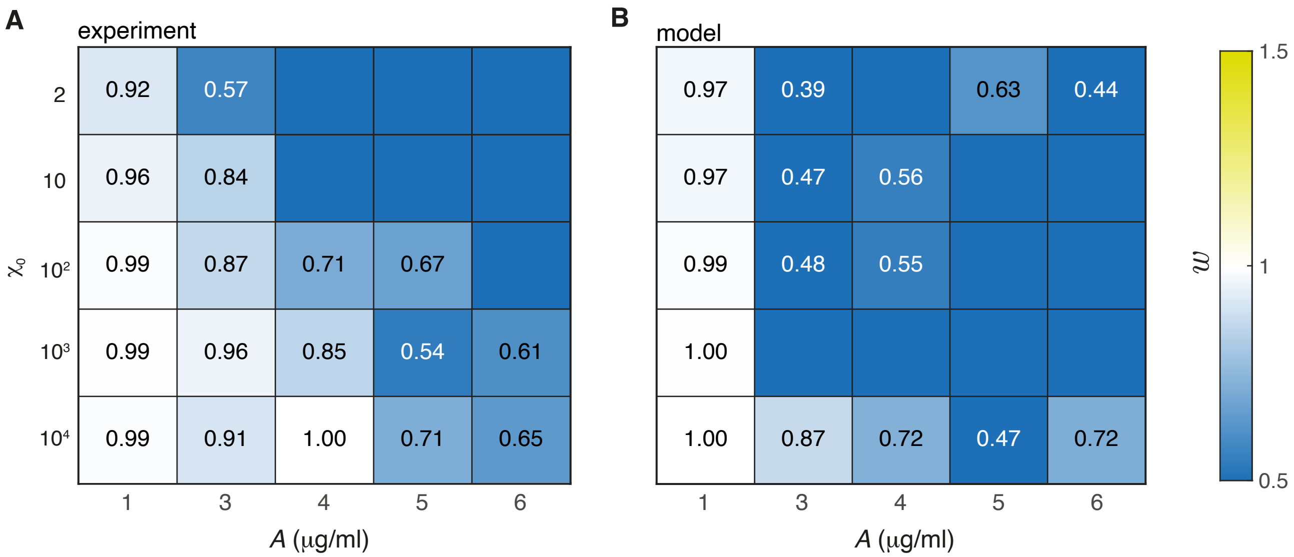


**Figure S6. Fitness of sensitive bacteria in respect to resistant bacteria in Fig. 2.** The average fitness of the periphery sensitive relative to the resistant strain, $w$, at various inoculation ratios, $\chi_0$. The value $w = 1$ signifies equal fitness and $w < 1$ ($w > 1$) that the sensitive strain is disadvantaged (advantaged) relative to the resistant. $N = 2$ for all entries. **A:** Experimental results for $OD_{600}$=0.3 on LB-agar with different Cm levels and incubated for 18 h. **B:** Modeling results (data from Fig. 2 in the main text) with $N = 1$. Empty entries with no value signifies divergence of $w$, when $\phi(18\,\mathrm{h}) = 0$ (i.e., sensitive did not reach the edge).

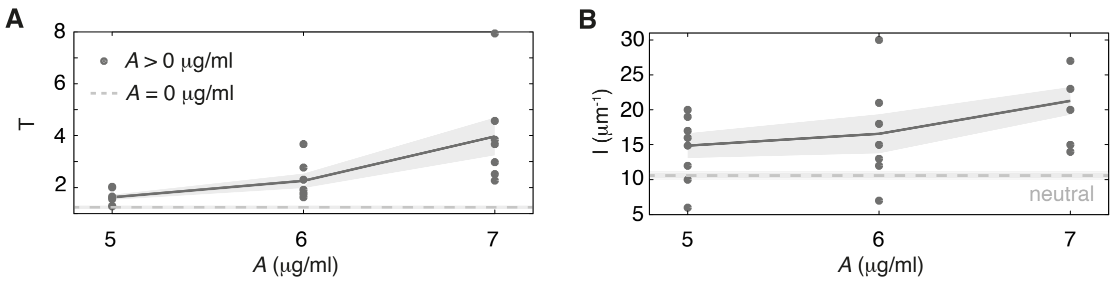


**Figure S7. Entanglements of colliding fronts versus Cm concentrations. A-B:** Tortuosity, $T$ (A), and intermixing index, $I$ (B), of the colliding fronts in Fig. 1B in the main text. Dots are experimental replicates of colliding fronts, full line are the average values, punctuated lines are $[\mathrm{Cm}] = 0\ \mu\mathrm{g/ml}$ and the shaded regions correspond ±SEM.

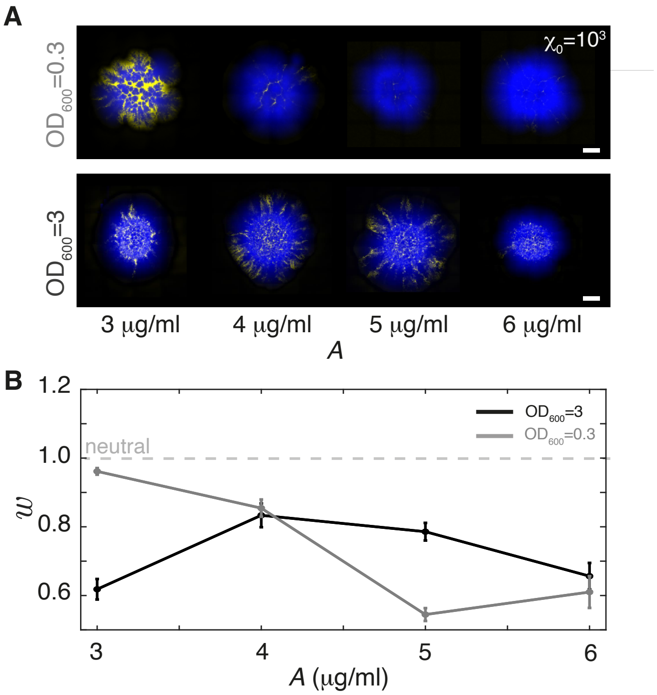


**Figure S8**. **Proximity favors the rescue of sensitive bacteria. A:** Pseudo-colored confocal laser-scanning images of surface-attached co-culture colonies of resistant (blue) and sensitive (yellow) at $\chi_0 = 10^3$ and densities of either $OD_{600}$=0.3 (similar to Fig. 2) or $OD_{600}$=3 on LB-agar plates with different Cm concentrations and incubated for 18 h. The scale bars correspond to 1 mm. **B:** The average fitness of the sensitive relative to the resistant strain, $w$, at various inoculation densities, $OD_{600}$ (A and legends). The horizontal dashed line corresponds to equal fitness ($w = 1$) and the shaded area corresponds to ±SEM and $N = 2$ for all entries (except for 3 µg/ml and $OD_{600}$=0.3, where $N = 4$).
